# Structural basis for immune cell binding of *Fusobacterium nucleatum* via the trimeric autotransporter adhesin CbpF

**DOI:** 10.1101/2024.09.17.613310

**Authors:** Gian Luca Marongiu, Uwe Fink, Andreas Oder, Jens Peter von Kries, Daniel Roderer

## Abstract

*Fusobacterium nucleatum* (Fn), a commensal in the human oral cavity, is overrepresented in the colon microbiota of colorectal cancer (CRC) patients and is linked to tumor chemoresistance, metastasis, and a poor therapeutic prognosis. Fn produces numerous adhesins that mediate tumor colonization and downregulation of the host’s anti-tumor immune response. One of these, the trimeric autotransporter adhesin (TAA) CEACAM binding protein of *Fusobacterium* (CbpF), targets CEACAM1 on T-cells and has been associated with immune evasion of Fn-colonized tumors. Whereas the role of CEACAM1 in homophilic and heterophilic cell interactions and immune evasion is well described, the mechanistic details of its interaction with fusobacterial CbpF remain unknown due to the lack of a high-resolution structure of the adhesin-receptor complex.

Here, we present two structures of CbpF alone and in complex with CEACAM1, obtained by cryogenic electron microscopy and single particle analysis. They reveal that CbpF forms a stable homotrimeric complex whose N-terminal part of the extracellular domain comprises a 64 Å long β roll domain with a unique lateral loop extension. CEACAM1 binds to this loop via its N-terminal IgV-like domain with high affinity with a nanomolar dissociation constant, as obtained by surface plasmon resonance. This study provides the first structural description of a fusobacterial TAA, illustrates a yet undescribed CEACAM1 binding mode, and paves the way for rational drug design targeting Fn in CRC.

## Introduction

*Fusobacterium nucleatum* (Fn), a commensal of the human oral cavity associated with various forms of periodontal disease, has been found to be overrepresented among the gut microbiota of colorectal cancer (CRC) patients (Dai et al., 2018). Although expressing no known toxins, Fn has been found to exacerbate cancer progression; Fn presence in CRC patients has been associated with poor prognosis, chemoresistance and metastasis (Mima et al., 2016, 2016; Xu et al., 2021).

While some mechanistic evidence is still missing, some adhesins have been described that exert secondary functions in addition to adhesion. The filamentous FadA binds to E-cadherin which is often overexpressed on cancer cells, activating the β-catenin/Wnt-signaling pathway leading to hyperproliferation of the respective cells (Guo et al., 2020; Rubinstein et al., 2013). The bifunctional adhesin Fap2 has been demonstrated to adhere to cancer cell-specific Gal-GalNAc-moieties while also deactivating tumor infiltrating lymphocytes (TILs) through the immune cell receptor TIGIT (Abed et al., 2016; Gur et al., 2015). Fap2 is a type Va autotransporter adhesin (Umana et al., 2019) and forms a ∼45 nm long β-helical rod-shaped extracellular domain with the proposed interactions sites for its two receptors at the membrane-distal tip of the rod (Schöpf et al., 2024).

In addition to the Fap2-mediated inhibition of immune cells via TIGIT interaction, another TIL inactivation pathway of Fn has been described that involves interaction of Fn with the carcinoembryonic antigen-related cell adhesion molecule 1 (CEACAM1) (Gur et al., 2019). Recently, the trimeric autotransporter adhesin (TAA) CEACAM binding protein of *Fusobacterium* (CbpF) was identified that targets CEACAM1 on T-cells and natural killer cells. In contrast to Fap2, CbpF is a homotrimeric type Vc autotransporter protein, and all three protomers together form a trimeric passenger domain, which remains exposed on the cell surface after expression, and a C-terminal β-barrel domain, serving as a membrane anchor (Brewer et al., 2019; Shhadeh et al., 2021).

CEACAM1, also known as cluster of differentiation 66a (CD66a), is the primordial member of the CEACAM family of glycosylated immunoglobulin (Ig) molecules (Kim et al., 2019). The extracellular domain (ECD) of CEACAM1 comprises an N-terminal, membrane-distal immunoglobulin variable region-like (IgV-like) domain and three membrane-proximal immunoglobulin constant region-like (IgC2-like) domains IgC2-like A1, IgC2-like B, and IgC2-like A2. While the IgC-like domains are heavily glycosylated, the IgV-like domain that is involved in interaction with CbpF (Brewer et al., 2019) is less glycosylated (Gray-Owen and Blumberg, 2006).

In physiological contexts, CEACAM1 is involved in cell adhesion and signaling. Its expression is constitutive and tightly regulated, and it is found on selected epithelial, endothelial, lymphoid, and myeloid cells (Prall et al., 1996; Odin and Obrink, 1987; Gray-Owen and Blumberg, 2006; Kammerer et al., 1998). CEACAM1 is involved in *trans*-homophilic and heterophilic interactions that mediate adhesion across epithelial cells and prevent self-damaging autoimmune responses between epithelial and immune cells (Gray-Owen and Blumberg, 2006; Kim et al., 2019). In pathological contexts, CEACAM1 is strongly expressed on various cancer cells, such as melanoma (Ullrich et al., 2015), colorectal (Neupane et al., 2021), lung (Zhou et al., 2013) and pancreatic cancer (Gebauer et al., 2014). This is assumed to promote a pro-tumorigenic tumor microenvironment by facilitating the evasion of cancer cells from immune cell-mediated killing, because trans-interaction of CEACAMs between epithelial and immune cells has been shown to inhibit T, B, and natural killer cells (Gray-Owen and Blumberg, 2006). In a seemingly synergistic manner, CbpF-mediated CEACAM1 binding down-regulates immune cells in the vicinity of Fn, thereby aiding immune evasion (Galaski et al., 2021).

These findings together explain why Fn is a great risk factor for cancer patients, making CbpF and other Fn adhesins important drug targets. The CbpF-CEACAM1 interaction has been analyzed by mutation studies, which have revealed a set of important interface residues in CEACAM1 (Brewer et al., 2019). However, no high-resolution structure of CbpF and its complex with CEACAM1 is available, which prevents us from understanding the detailed molecular mechanism of their interaction. Here, we present high-resolution structures of CbpF alone and in complex with the CEACAM1-ECD. Cryogenic electron microscopy (cryo-EM) and single particle analysis (SPA) revealed that the N-terminal part of the CbpF extracellular domain (ECD) comprises a threefold symmetric, parallel β roll. The CEACAM1 N-terminal IgV-like domain binds to a unique loop protruding from β-sheet stack of the β roll. Furthermore, we demonstrate the CbpF interaction *in cellulo* with purified adhesin on cells and report submicromolar affinity *in vitro*, determined by surface plasmon resonance (SPR). Collectively, this work provides the prerequisite for structure-based drug design through a detailed understanding of the CbpF/CEACAM1 complex assembly.

## Results

### Structure of recombinantly purified CbpF reveals a homotrimeric complex with a β roll

To understand the assembly mechanism of the trimeric type Vc autotransporter protein CbpF, we initially aimed to determine its structure at near atomic resolution. For this, we used a codon-optimized version of *F. nucleatum* ATCC2586 FN1499 that codes for CbpF and cloned the sequence corresponding to the mature protein after signal peptide cleavage (residues S24 – K479) in pASK-IBA2C in frame with an N-terminal *Escherichia coli* OmpA signal peptide and a C-terminal StrepII-tag. After recombinant expression, we extracted the protein from the membranes adapting an established protocol for the purification of *Yersinia enterocolitica* YadA (Chauhan et al., 2019) and purified it via affinity chromatography to Strep-Tactin resin. In agreement with previous findings (Brewer et al., 2019; Shhadeh et al., 2021), a prominent band in SDS-PAGE above 150 kDa indicates the formation of a stable trimer (SI Fig. 1A). We then reconstituted CbpF in nanodiscs composed of DMPC and the MSP1D1 scaffold protein. The SDS- and heat-resistant trimeric form of CbpF remained the predominant species, as judged by SDS-PAGE (SI Fig. 1B,C), indicating that nanodisc incorporation added further to stabilization of the complex. Indeed, we observed several rod-shaped protrusions adjacent to nanodiscs in the reconstituted CbpF preparation in negative staining electron micrographs that are in agreement with the ECD of a trimeric autotransporter (SI Fig.1D).

We next vitrified nanodisc-embedded CbpF for cryo-EM and SPA, and obtained a density map with a final resolution of 3.8 Å (SI Fig. 2). Using and adjusting a model predicted by AlphaFold2 complex (Bryant et al., 2022) revealed that the N-terminal part of the ECD (residues A25 – G274) fitted in the density (Fig. 1A). The rest of the ECD after G274, and the C-terminal transmembrane barrel (predicted residues V378 – L431) were not part of the highly resolved structure (Fig. 1B). The model of the region after G274 was predicted to be without secondary structure and thereby likely highly flexible, matching the disappearance of signal in the density map. Within the N-terminal part of the ECD, V28 – N216 form a β roll domain of ∼64 Å length and ∼44 Å diameter at its widest part, comprising 14 parallel β-strands per protomer (Fig. 1A, C). The last β-strand (R212 – N216) is swapped from the neighboring protomer into the stack of β-strands. C-terminal of the β roll, intertwining loops and short α-helices (residues D219 – G274) form a ∼54 Å long extension (Fig. 1D) ahead of the unstructured, not resolved region. This region might function as a spacer to extend the distance of the N-terminal, receptor-binding part of CbpF from the Fn outer membrane, similar as observed in other autotransporter adhesins (Melia et al., 2021). The autotransporter region of CbpF with the nanodisc became evident in several low resolved 2D classes in the data set (SI Fig. 3A). 3D reconstruction and refinement of this small subset of 21,632 particles resulted in an 8.3 Å density map, in which the predicted autotransporter region and its adjacent α-helix linkers fitted, but also here the most flexible part was not resolved (SI Fig. 3B-D). Therefore, we expect that the relative position of the N-terminal ECD with respect to the C-terminal autotransporter of CbpF is highly variable, allowing flexible attachment to target cells via CEACAM1 in functional analogy to a fishing rod and line.

**Figure 1:**
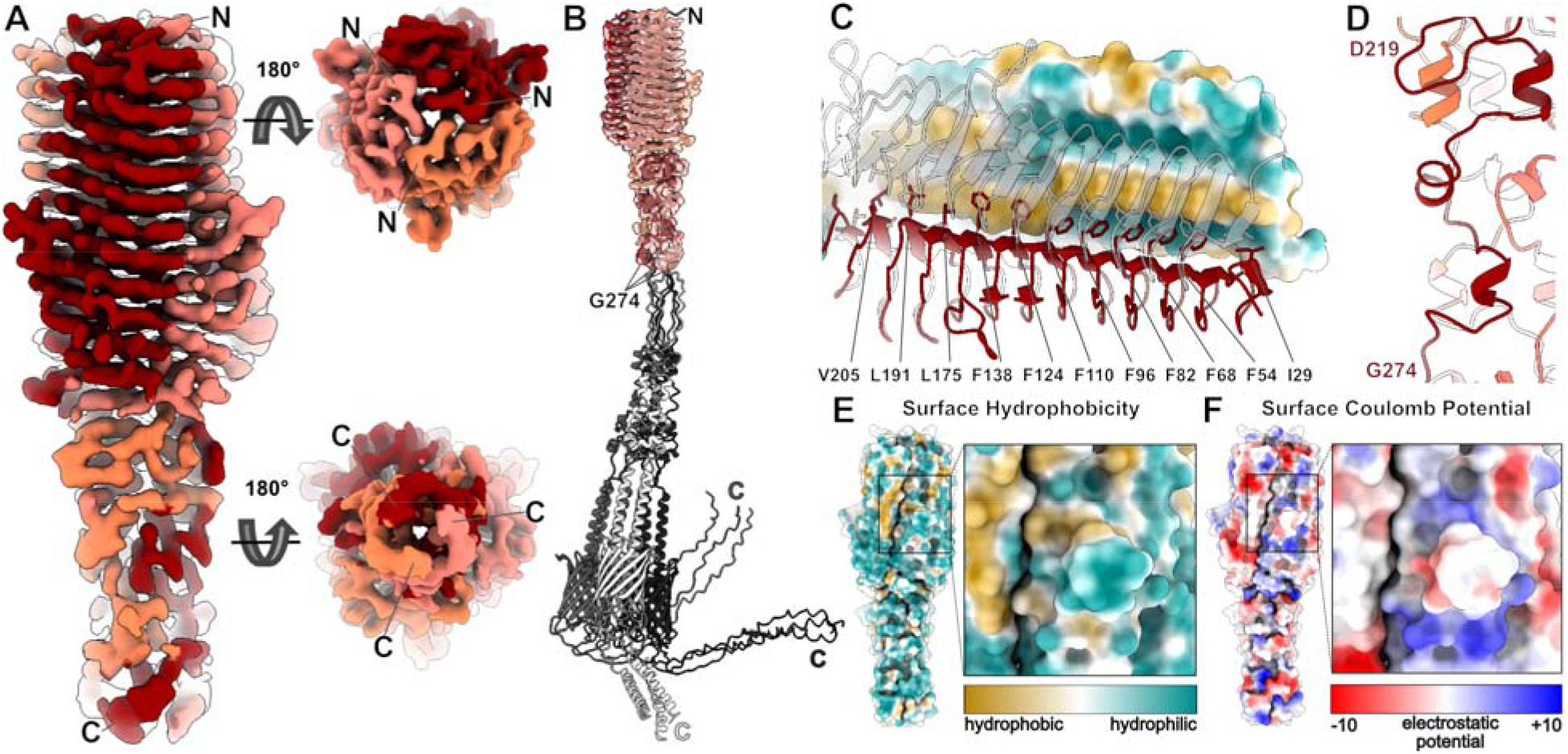
Structure of the *F. nucleatum* type Vc autotransporter adhesin CbpF. A: Cryo-EM density of CbpF at 3.8 Å resolution. The map is colored in red tones corresponding to the three protomers. N-and C-termini of the resolved parts are indicated. B: Density map colored as in A with the three highest scoring AlphaFold2 models of the CbpF homotrimer fitted. The non-resolved parts after G274 are colored in gray tones. Note the unstructured regions following after G274. C: Inward-facing hydrophobic F, I, L, and V residues form the hydrophobic core of the β roll and hold together the protomers within the homotrimeric passenger domain. D: Immediately C-terminal of the β roll, short α-helices and loops of the three CbpF protomers wrap around each other. E,F: Surface hydrophobicity (E) and Coulomb potential (F) of the CbpF β roll and the following α-helical domain. A loop protruding from the β-sheet stack is highlighted and shown in zoomed view.

To deduce the potential receptor binding site of CEACAM1 to CbpF, we inspected the surface hydrophobicity and charge distribution of the CbpF ECD. This revealed a hydrophobic strip adjacent to the contact sites of two protomers in the center of the β roll (Fig. 1E). Directly next to this strip, a loop that is made up of residues D142 - Y149 protrudes out of the stack of β-sheets, whose outward-facing surface is hydrophilic whereas the upper part of its inward-facing surface is hydrophobic (Fig. 1E). One of the loop’s sides is negatively charged, whereas its surrounding is mostly positively charged at neutral pH (Fig. 1F). Therefore, we proposed that the loop is the possible CEACAM1 binding site and subjected it to a detailed inspection.

### A unique loop protrudes from the CbpF β roll that is only conserved in *Fusobacteria*

We then analyzed the conservation of the loop region (D142 - Y149) in CbpF-homologs of other *Fusobacteria*. We found close homologs in the closely related Fn ATCC23726 (Fusoportal gene 4), *F. vincentii* ATCC49256 (FNV1729), for which CEACAM1 binding is known (Brewer et al., 2019), *F. polymorphum* (FNP1391), and *F. necrophorum* (1_1_36S; SI Fig. 4A), with overall sequence identities between 94 and 42% to CbpF of Fn ATCC25586 (SI Fig. 4B). The sequence of the loop is fully conserved in Fn ATCC23726 and conserved with one Q-to-S point mutation in *F. vincentii*. In contrast, it is not conserved in the other two *Fusobacteria*, where longer insertions are evident (Fig. 2A). Therefore, these two might not bind to CEACAM1, as observed for two clinical Fn isolates 2B24 and 2B28 and *F. periodonticum* (Brewer et al., 2019), of which the latter one (sequence 2_1_31 (Sanders et al., 2018)) has no conserved CbpF sequence. Therefore, we deduce that only select *Fusobacteria* are able to attach to CEACAM1 via a CbpF-like autotransporter.

**Figure 2:**
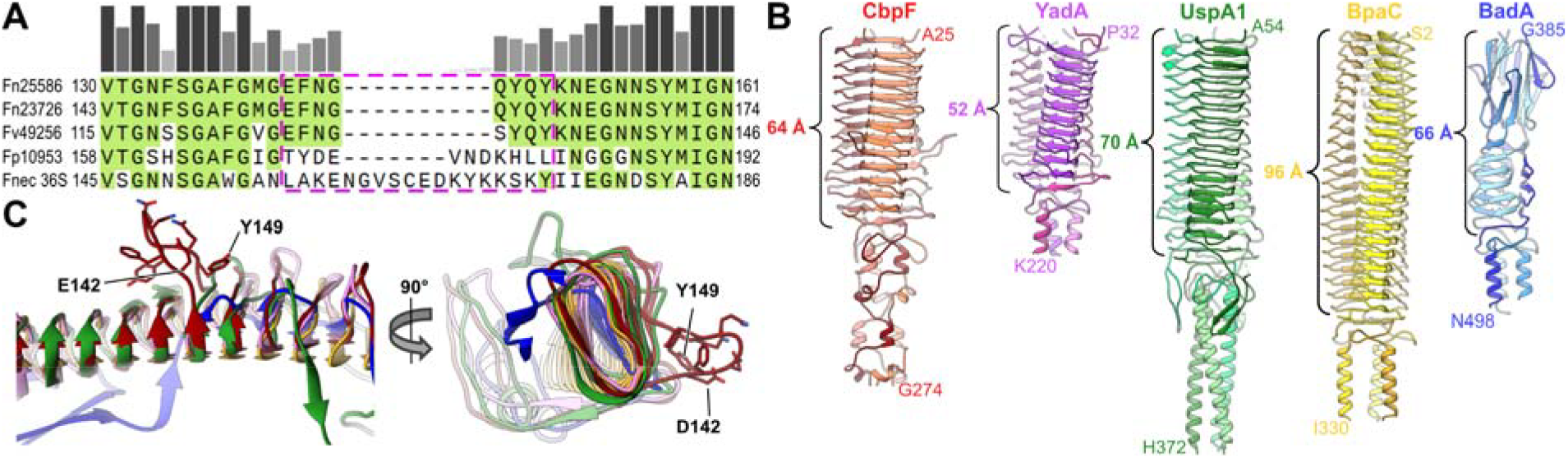
Conservation of loop and comparison of CbpF structure with other Type Vc autotransporter adhesins. A: Alignment of CbpF sequences from different *Fusobacteria* (Fn ATCC25586 – the present structure, Fn ATCC23726, *F. vincentii* (Fv) ATCC49256, *F. polymorphum* (Fp) ATCC10953, and *F. necrophorum* 1_1_36S (Fnec)). The section shows the region around the loop E142 – Y149 in Fn ATCC25586, which is highlighted with pink dashed frame. B: Comparison of the passenger domains of Fn CbpF with *Yersinia enterocolitica* YadA (PDB 1P9H), *Moraxella catarrhalis* UspA1 (PDB 3PR7 and 3NTN), *Burkholderia pseudomallei* BpaC (PDB 7O23), and *Bartonella henselae* BadA (PDB 3D9X). The lengths of the β roll domains (β-sheet stack and Trp ring domain for BadA) are indicated. C: Structure alignment of one protomer of the five adhesins shown in B, with residues 142 - 149 from CbpF that form a loop protrusion outside the β roll highlighted. None of the other adhesins has a similar loop. Note that UspA1 is also a CEACAM1 binding protein.

To assess whether the loop or a similar protrusion from the β roll of autotransporter adhesins is structurally conserved, we compared CbpF with similar type Vc autotransporter adhesins with available structures in the protein data bank (PDB). We found adhesins with similar ECDs from *Yersinia enterocolitica* (YadA) (Nummelin et al., 2004), *Moraxella catarrhalis* (UspA1) (Agnew et al., 2011), *Burkholderia pseudomallei* (BpaC) (Kiessling et al., 2022), and, with a different organization of the β-sheet stack, *Bartonella henselae* (BadA) (Szczesny et al., 2008). The length of the β roll, which is 64 Å in CbpF, varied between 52 Å for YadA (11 β-sheets in stack) and 96 Å for BpaC (21 β-sheets), whereas their diameters appeared nearly identical (Fig. 2B). Structural alignment of the β-sheet domains revealed that none of the adhesins except CbpF has a similarly sided and shaped loop protruding from the stack (Fig. 2C). Even in UspA1, which also binds to CEACAM1 via another motif (Conners et al., 2008), not a long loop but only a very short protrusion made up from 2 residues is present, as it is in YadA, and nothing at all is present in the other two adhesins (Fig. 2C). Therefore, we conclude that the loop aside the β roll is an exclusive feature of fusobacterial CbpF and contains a hitherto undescribed CEACAM1 binding motif.

### CbpF binds to CEACAM1 with nanomolar affinity *in vitro*

Next, we tested binding activity of the recombinantly produced CbpF to immune cells. For this, we fluorescently labeled nanodisc-embedded CbpF and incubated it with the T-cell line Jurkat. After 2 h of incubation, weak fluorescence signals that originate from CbpF were evident on the surface and also already within Jurkat cells, the latter due to endocytosis. Jurkat cells that have been incubated with nanodiscs labeled in the same way did not show comparable fluorescence signals (Fig. 3A,B). Similar observations with an even more pronounced CbpF binding were made for human embryonic kidney (HEK293) cells (SI Fig. 5A), which constitutively express CEACAM1, as described previously (Dery et al., 2011) and annotated in the human proteome atlas (Uhlén et al., 2015). This demonstrates that recombinant CbpF is functional, and that CbpF alone is sufficient to attach to human cells that display CEACAM1.

**Figure 3:**
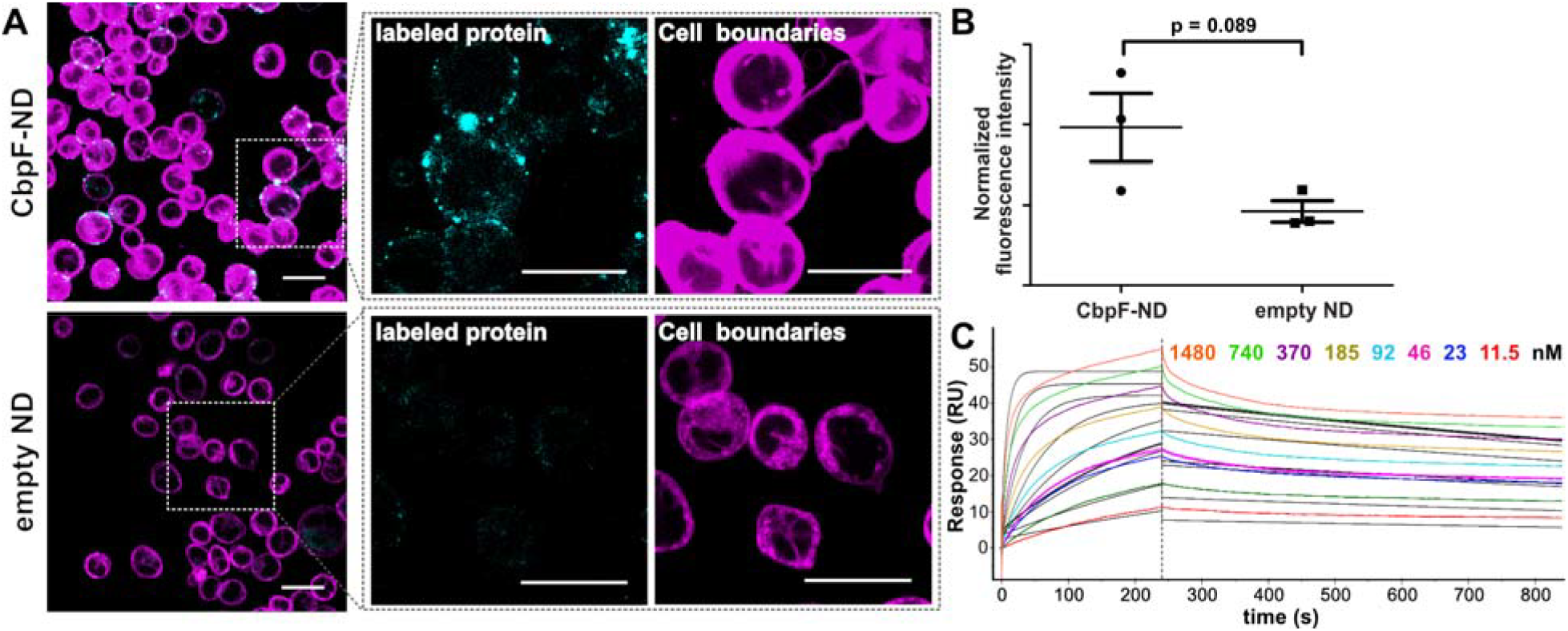
Binding and affinity of *F. nucleatum* CbpF to human CEACAM1. A: Exemplary images of Jurkat cells that bind nanodisc-reconstituted CbpF (top) or empty MSP1D1 nanodiscs (lower panels). Cyan channel: protein labeled with AF 488, violet channel: cell boundaries labeled with CellMask™ Deep Red Plasma Membrane Stain. Scale bar: 250 µm. B: Quantification of cell binding as in A. At least 39 regions of interest (ROIs) were analyzed per replicate. C: Surface plasmon resonance sensorgram showing binding of CbpF to immobilized CEACAM1 ECD. The concentrations of CbpF are indicated. The black solid lines show a global fit of the binding curves according to a 1:1 binding model and reveal a k_on_ of 8.13 ± 0.01 × 10^4^ M^-1^ s^-1^ and k_off_ of 5.1 ± 0.02 × 10^−4^ s^-1^, resulting in a K_D_ of 6.2 nM.

We next determined the yet unknown affinity of CbpF to CEACAM1 using surface plasmon resonance (SPR). For this, we immobilized CEACAM1 ECD (residues 35-428, as obtained from ACRO Biosystems) and bound CbpF in solution. With a 1:1 binding model and global fit of 8 curves with different CbpF concentrations, we obtained a dissociation constant (K_D_) of ∼6 nM, with k_on_ of 8.13 ± 0.01 × 10^4^ M^-1^ s^-1^ and k_off_ of 5.1 ± 0.02 × 10^−4^ s^-1^ (Fig. 3C, SI Fig. 5B). With this, CbpF has a two orders of magnitude higher affinity to its respective immune cell receptor than the other known fusobacterial interaction with the immune system, i.e., the adhesin Fap2 with TIGIT (K_D_ of ∼600 nM (Schöpf et al., 2024)). Evaluating the steady-state signals of the binding curves, we obtained a slightly weaker affinity with a K_D_ of 75 ± 14 nM (SI Fig. 5C); yet, this still means an order of magnitude higher affinity compared to Fap2 and TIGIT. This clearly underlines the importance of CbpF in immune evasion of Fn and of tumors colonized by Fn.

Using the 1:1 binding model, the fit of the association and dissociation curves is not exact (SI Fig. 5B), with deviations especially in the association curves. Considering that CbpF is a trimer and that association with one CEACAM1 molecule might influence binding of further ones, we tested a heterogeneous ligand binding model. We obtained two K_D_ values of 2.5 and 47 nM, respectively, (SI Fig. 5D), which are in the same range as the value obtained with a 1:1 model and they do not suggest a great impairment of binding further CEACAM1 after the first one is bound to the CbpF trimer.

### CbpF binds to the N-terminal domain of CEACAM1 via the β roll protruding loop

Although CbpF and CEACAM1 form a stable complex and the structure of significant parts of CEACAM1 as well as CbpF–like type Vc proteins from other bacteria are known, structure prediction of the complex did not reveal interaction sites. Neither hetero-tetrameric (CbpF trimer with 1 CEACAM1) nor hetero-hexameric (CbpF trimer with 3 CEACAM1) complexes were reproducibly predicted with high confidence by AlphaFold2 multimer (SI Fig. 6). This indicates a yet unknown interaction mechanism, likely involving the identified loop as described in Fig. 2.

We therefore formed the complex of nanodisc-reconstituted CbpF and CEACAM1-ECD (residues 35-428, ACRO Biosystems) and removed excess of CEACAM1 by preparative size exclusion chromatography (SEC, SI Fig. 7A). Due to glycosylation, CEACAM1 appeared as a broad and faint band on SDS-PAGE (SI Fig. 7B), which is not present in SDS-PAGE of nanodisc-embedded CbpF alone. We then vitrified the SEC fraction corresponding to the main peak (SI Fig. 8A) and resolved the structure of the complex to 2.7 Å, applying C3 symmetry (Fig. 4A, SI Fig. 8B-H). The structure revealed a hetero-hexameric complex in which one CEACAM1 is bound per CbpF protomer, with the N-terminal IgV-like domain of CEACAM1 and the region around the β roll protruding loop of CbpF forming the interface (Fig. 4A,B). Each CEACAM1 protrudes perpendicularly from the central CbpF trimer and sits atop the β roll protruding loop of CbpF. Whereas the IgV-like domain has been confirmed to interact with CbpF previously (Brewer et al., 2019) and is also responsible for CEACAM1 interaction with other adhesins (Conners et al., 2008; van Sorge et al., 2021), the CbpF part of this interface has no homologies in other complexes. Four residues of the eight-residue long loop 142-149, i.e., E142, N144, Q146, and Q148, form H-bonds with side chains of T90, Q78 (Q44 in mature chain (Brewer et al., 2019)), and Y68 of CEACAM1. In addition, there are two further H-bonds three β strands “above” in the stack formed between H92 and N99 of CbpF and the backbone of S127 and D128 of CEACAM1 (Fig. 4C). Importantly, H92 originates from the neighboring CbpF protomer, and both together form one CEACAM1 interface. This demonstrates that the trimerization of CbpF is not only relevant for its stability, but also for its receptor-binding functionality.

**Figure 4:**
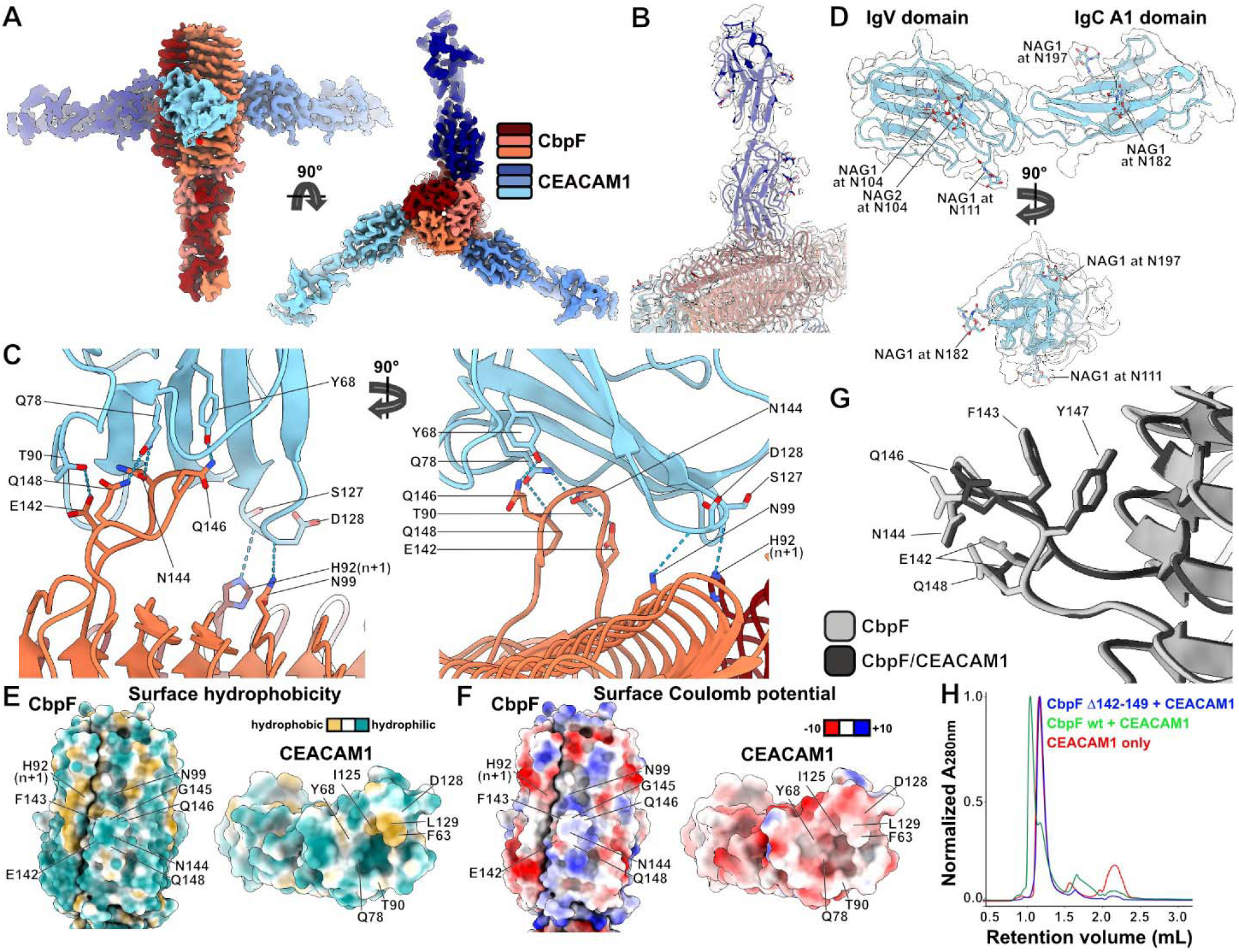
Structure of the CbpF-CEACAM1 complex. A: Cryo-EM density of the complex at 2.7 Å resolution. The map is colored in red tones corresponding to the three CbpF protomers and in blue tones corresponding to three CEACAM1 molecules, in which the IgV-like and IgC2-like A1 domains are resolved. B: Model-to-map fit focused on one CEACAM1 bound to CbpF. Color code of the model as the maps in A. The density map is shown in transparent gray. C: Model of the CbpF/CEACAM1 interface. Intermolecular H-bonds as obtained in ChimeraX are shown as dashed lines. Note that H92 from CbpF protomer n+1 (dark red) contributes to the interface of CbpF protomer n (coral). D: Cryo-EM density map of one CEACAM1 at 3.0 Å resolution (transparent gray), obtained by shifting the center of reconstruction of the complex map, with the fitted model including glycan moieties. E, F: Surface hydrophobicity (E) and Coulomb potential of the interacting sites of CbpF (left) and CEACAM1 (right) from the complex. The views show the front views of the interacting sites. G: Overlay of CEACAM1-interacting loops of CbpF in the complex (black) with unbound CbpF (gray). H: SEC profiles on a Superdex 200 3.2/300 column that show that CbpF wt (33 µg total protein amount, 1:1 ratio) forms a complex with CEACAM1, as evident by a peak shift to earlier retention volume, whereas CbpF Δ142-149 mutant (50 µg total protein amount, 1:1 ratio) does not. Note that CEACAM1 co-elutes with the CbpF mutant at the same retention volume.

Whereas previous structural analysis by X-ray crystallography only resolved the N-terminal IgV-like domain of CEACAM1 (residues 35-142), our cryo-EM structure, after shifting the center of reconstruction towards one CEACAM1, also resolves the following IgC2-like A1 domain (res. 35-234 in total, SI Fig. 8I-N). Although the ECD of CEACAM1 is highly glycosylated and our data revealed several glycosylation sites on the IgV-like and IgC2-like A1 domains (Fig. 4D), no glycans are involved in interaction with CbpF. Instead, a tight interface is formed exclusively by the peptide moieties predominantly via H-bonds (Fig. 4C), whereas only two hydrophobic surface-exposed residues (F63 and L129 in CEACAM1) are buried in the complex (Fig. 4E). The rest of the CEACAM1 side of the interface appears slightly negatively charged, whereas at the CbpF side of the interface positive charges dominate (Fig. 4F). A hydrophobic stretch at the contact site of two protomers adjacent to H92 flanks the binding region, but is not involved in direct interactions with CEACAM1 residues (Fig. 4E). The same hydrophobic patch and also a positive charge around the binding site is evident in the structure of CbpF alone (Fig. 1E,F), and comparison of the CEACAM1 binding loop between CbpF and the complex reveals a nearly identical arrangement (Fig. 4G). This shows that the interface is already formed out in CbpF without the need for a conformational transition upon binding.

To test whether the CbpF/CEACAM1 complex can also form without the β roll protruding loop, we removed CbpF residues 142-149 by site-directed mutagenesis. We then expressed and purified the resulting CbpF Δ142-149 mutant, mixed it with CEACAM1 in a 1:1 molar ratio and checked complex formation via SEC. Clearly, there is no shift of the main peak towards lower retention volumes for CbpF Δ142-149/CEACAM1, as it is observed for wild type (wt) CbpF/CEACAM1 (Fig. 4H). Even at a higher CbpF Δ142-149 concentration than CbpF wt, there are no traces of a complex formed, clearly highlighting the importance of the loop.

Taken together, the CbpF/CEACAM1 complex structure, which is to our knowledge the first structure of a fusobacterial trimeric autotransporter adhesin-receptor complex, demonstrates a hitherto undescribed binding mode of bacterial adhesins to immune proteins. This mechanism appears to be specific to CbpF and similar adhesins of closely related *Fusobacteria*.

## Discussion

Our structure of the CbpF/CEACAM1 complex reveals intricate details of the interaction. CbpF binds to CEACAM1 via a protruding loop (D142 - Y149) and via H92 on the respective adjacent protomer chain. Comparison to similar TAAs render this a hitherto undescribed CEACAM1 binding motif. Even though CEACAM1 is highly glycosylated, no glycans are involved in CbpF/CEACAM1 complex assembly.

It has been demonstrated that CbpF/CEACAM1 complexation is dependent on its trimeric assembly, other than that of a similar TAA from *Moraxella catarrhalis*, UspA1. While a single UspA1 protomer is sufficient to bind to CEACAM1 (Hill et al., 2012), only trimeric CbpF binds to CEACAM1 (Brewer et al., 2019). Our findings demonstrate that not only the CbpF protomer chain with the protruding loop in close contact with the CEACAM1 dimerization site, but also H92 of the adjacent protomer contributes to binding. Consequently and in agreement with previous observations (Brewer et al., 2019), a single protomer chain would not form a complete CEACAM1-binding interface.

Previous site-directed mutagenesis studies showed that alteration of F63 and Q78 of CEACAM1 each abolishes CbpF/CEACAM1 binding (Brewer et al., 2019). Our CbpF/CEACAM1 complex structure shows that CEACAM1-F63 is in close proximity to H92 and F143, which probably form aromatic stacking interactions. Conserved Q78 of CEACAM1 forms H-bonds with N144 and Q148 of CbpF. This underlines the critical relevance of this central H-bond that coordinates two residues of CbpF.

Besides CbpF, other bacterial adhesins capable of binding to CEACAM1 have been described, such as the TAA UspA1 from *M. catarrhalis* (Fig. 2b), which has been shown to bind to CEACAMs (Hill et al., 2005). Although the β rolls of CbpF and UspA1 are structurally similar, CEACAM1 binding of UspA1 is not located in this region as in CbpF, but instead in a long α-helical coiled-coil stalk. UspA1/CEACAM1 binding is mediated by a coiled-coil motif (Conners et al., 2008), which lacks the protruding loops identified in CbpF. Another CEACAM-binding bacterial adhesin is the immunoglobulin-like adhesin β protein of *Streptococcus agalactiae*, which has been reported to bind to CEACAM1 with a novel Ig-like fold termed IgI3 (van Sorge et al., 2021). Although structurally very different, the CEACAM1-binding interfaces of CbpF and IgI3 share common features. Both employ a protruding hydrophilic loop with an adjacent hydrophobic stretch. In addition to TAA-like and Ig-like CEACAM-binding adhesins, β-barrel-shaped adhesins such as OMP P1 of *Haemophilus influenzae* (Tchoupa et al., 2015) and OPA of *Neisseriae* sp. (Kuespert et al., 2011; Robertson and Meyer, 1992) have also been reported. This shows that although the CEACAM1 binding motifs are very different, host cell binding via CEACAM1 is widespread in bacterial pathogens, which indicates that this interaction has evolved independently.

In physiological contexts, CEACAM1 forms trans-homodimers at the GFCC’ face (CEACAM1 β-strands G, F, C, and C’) via, among other residues, Y68 and Q78 (Gandhi et al., 2021), that are also critical for the interaction with CbpF via Q146, N144, and Q148. From the other CEACAM-binding adhesins described above, group B *Streptocoocus* IgI3 targets CEACAM1-Q78, while UspA1 interacts with both CEACAM1-Y68 and Q78. Also other known bacterial adhesins with the capacity to bind CEACAM1, i.e., AfaE of *Escherichia coli* (Anderson et al., 2004) and HopQ of *Helicobacter pylori* (Javaheri et al., 2016), target the CEACAM1 dimerization interface. This shows that despite the structural variability of bacterial CEACAM-interaction motifs, the CEACAM1 target site appears to be conserved.

In addition to homodimerization, CEACAM1 also forms dimers with CEACAM5 (Gandhi et al., 2021). Afa adhesins of *E. coli* have been shown to target the CEACAM family members CEACAM3, -5, and -6 in addition to CEACAM1 (Berger et al., 2004). In contrast, CbpF has been shown to bind only to CEACAM1 and CEACAM5 (CEA) (Brewer et al., 2019). The only two unique residues found in both CEACAM1 and CEACAM5, F63 and Q78, are both targeted by CbpF. Since CEACAM1 and CEACAM5 are the only members of the CEACAM family that share these residues, our results provide the structural basis for the selectivity of CbpF. This may allow Fn to avoid activation of CEACAM3-expressing neutrophils (Heinrich et al., 2016), which is in agreement with the overall functional scheme of adhesion combined with immunosuppression.

Our results demonstrate a strong CbpF interaction with CEACAM1 with nanomolar affinity. For the structurally similar TAAs YadA from *Yersinia* spp. and UspA1 from *M. catarrhalis*, no dissociation constants have been reported. Functionally, however, parallels are noteworthy. UspA1 from *M. catarrhalis*, targets fibronectin and laminin for adhesion (Brooks et al., 2008) and CEACAM for potential immune evasion.

Besides CbpF, Fn produces a multitude of other adhesins including the filamentous FadA (Rubinstein et al., 2013) and the bi-functional Fap2. The latter binds to the immune receptor TIGIT on tumor invading NK cells (Gur et al., 2015) and to the glycan Gal-GalNAc abundant on colon cancer cells (Abed et al., 2016). Thus, analogous to CbpF, Fap2 exerts adhesive and immunosuppressive functions (Fig. 5).

**Figure 5:**
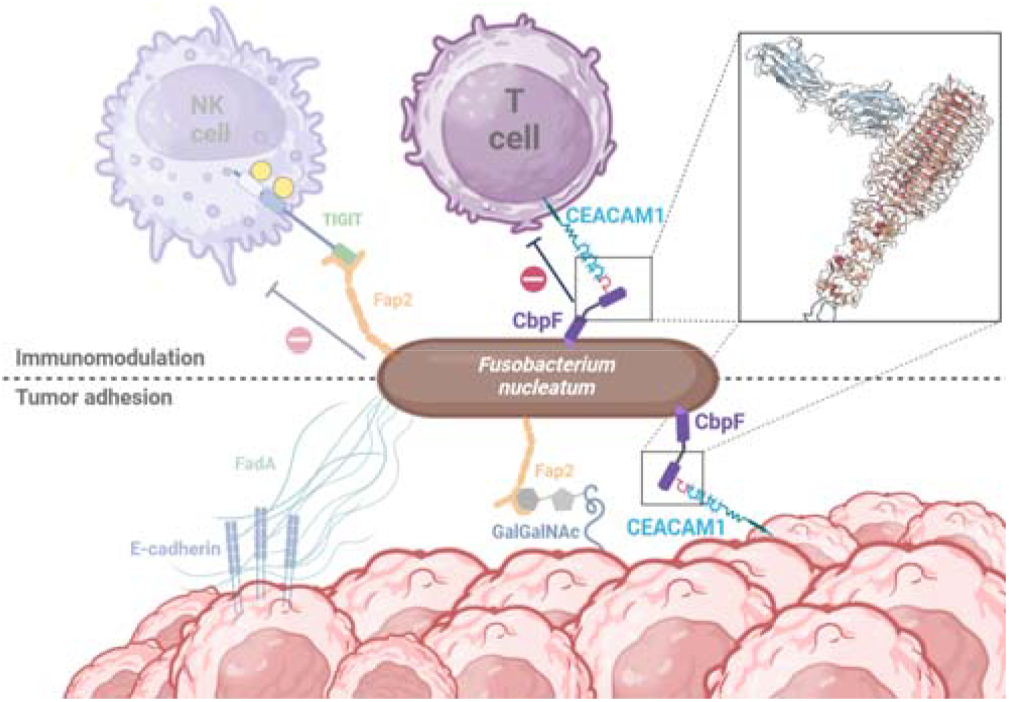
Model of CEACAM1 mediated adhesion of *F. nucleatum* to immune and cancer cells. *F. nucleatum* binds to and down-regulates the activity of T-cells via CbpF, while the adhesins FadA and Fap2 mediate interaction with CRC cells in tumors. In addition, CbpF might also be able to form a ternary complex with CEACAM1 on T-cells and on cancer cells, facilitating both immune evasion and tumor colonization.

Taken together, our findings provide the structural basis for understanding the unique interaction between CbpF and CEACAM1. The high binding affinity of CbpF/CEACAM1 suggests a role of CbpF not only in immune suppression, but possibly also in Fn tumor colonization. Additionally, the binding motif is distinct and selective, rendering this interaction a promising drug target.

## Methods

### Cloning of *F. nucleatum* CbpF expression plasmids

The DNA sequence corresponding to CbpF from *F. nucleatum* ATCC25586 (FN1499) was purchased as codon-optimized gene from GenScript and cloned in the expression plasmid pASK-IBA2C (IBA Lifesciences) in frame with an N-terminal *Escherichia coli* OmpA signal peptide and a C-terminal StrepII-tag, resulting in pASK-IBA2C-CbpF. The expression plasmid for CbpF mutant Δ142-149 was prepared from pASK-IBA2C-CbpF by removing the DNA sequence that codes for the loop using KLD mutagenesis (New England Biolabs). All sequences were verified by Sanger sequencing (LGC Genomics). *E. coli* TOP10 (Thermo Fisher Scientific (TFS)) was used for cloning.

### Expression and purification of CbpF

E. coli BL21 C43 was transformed with pASK-IBA2C-CbpF or pASK-IBA2C-CbpF Δ142-149 and 50 mL Luria Bertani (LB) medium with 34 mg/L chloramphenicol were inoculated with a single colony, followed by overnight incubation at 180 rpm shaking and 37 °C to obtain a starter culture. 4 L of LB medium with 34 mg/L chloramphenicol were then inoculated with the starter culture and incubated at 37 °C until a OD_600nm_ of 0.8. Expression was then induced by adding 100 ng/mL anhydrotetracycline (Merck), and incubation was continued overnight at 22 °C.

Cells were harvested (Beckman F10 BCL, 20 min at 6000 rpm and 4 °C), resuspend in PBS pH 7.4, and lysed by three passages through a LM10 microfluidizer. 1 mM PMSF and 1250 units benzonase (Mobitec, Cat. No. GE-NUC10700-01) were added to the lysate, which was then cleared by centrifugation (20 min at 8000 rpm and 4 °C). The supernatant was then centrifuged for 1 h at 38000 rpm and 4 °C in a TI45 rotor (Beckman) to pellet the membranes. The membrane pellet was resuspended in extraction buffer (25mM MOPS-NaOH, 150mM NaCl, 50mM EDTA, 10mM DTT, 0.1 mg/ml lysozyme, 1% lauroylsarcosine (Merck, Cat. No. L5777), pH 7.0) and resuspended using a Dounce homogenizer, followed by 2 h of incubation at 23 °C to solubilize CbpF. Ultracentrifugation (1 h, 38000 rpm at 20 °C in a TI 45 rotor) followed, and the supernatant was loaded on a 5 mL large self-casted StrepTactin Superflow High capacity (IBA Lifesciences) column equilibrated with buffer A (25mM MOPS-NaOH, 300 mM NaCl, 0.8% lauroylsarcosine pH 7.0). CbpF was eluted with 5 mM desthiobiotin in buffer A and analyzed via SDS-PAGE. Fractions that contained CbpF were pooled and loaded on a Superdex 200 increase 10/300 column (Cytiva) equilibrated in buffer A. Fractions of 0.4 mL were analyzed via SDS-PAGE and CbpF-containing fractions were pooled and used for nanodisc reconstitution or SPR.

### Nanodisc reconstitution of CbpF

1 mg CbpF in buffer A was mixed with nanodisc scaffold protein MSP1D1 in buffer A and DMPC solubilized in buffer A with 10% lauroylsarcosine. The molar ratio of CbpF, MSP1A1, and DMPC was 1:20:1600. The mixture was incubated for 1h at 23 °C, then the detergent was removed by three subsequent applications to 300 mg biobeads SM2 (Biorad) for 2 h each at 23 °C. The supernatant was loaded on a Superdex 200 increase 10/300 column equilibrated in SEC buffer (25mM MOPS-NaOH, 300 mM NaCl pH7.0). The fraction with the highest concentration as judged by SDS-PAGE was used for cryo-EM and complex formation with CEACAM1.

### Negative stain electron microscopy

4 µL of Nanodisc-embedded CbpF in reconstitution buffer was loaded on a freshly glow-discharged 300-mesh carbon-coated copper grid. After an incubation of 1 min, the sample was blotted with Whatman filter paper, washed by a drop of Milli-Q water with subsequent blotting and stained twice with 2% uranyl acetate solution for 1 min. The staining solution was removed with Whatman filter paper and grids were air-dried for at least 5 min at room temperature. Grids were imaged on a Talos L120 TEM (TFS) using a Ceta-16M CCD detector.

### Cell binding assay of CbpF

For CbpF/CEACAM1 binding analysis to Jurkat cells (ATCC), cells were cultured in RPMI 1640 medium (PAN-Biotech, Cat. No. 4251184102738) supplemented with 10% (v/v) fetal bovine serum (PAN-Biotech, Cat. No. 4251184108365). Growth medium was aspirated after pelleting 1 × 10^8^ cells at 70 x g. Cells were resuspended in 1 mL DPBS (Pan Biotech, Cat. No. 4251184103971), split into two 1.5 mL tubes, washed with 500 µL DPBS, resuspended and pelleted at 1020 x g for 2 minutes. Cells were incubated with either 1 µM AF 488 NHS-ester (Lumiprobe, Cat. No. 11820) conjugated CbpF nanodiscs or AF 488 NHS-ester conjugated nanodiscs at 37 °C and 150 rpm for 2 hours. Fixation was carried out with 200 µL 4% (v/v) paraformaldehyde (Roth, Cat. No. 0335.1) at room temperature for 20 minutes. Cell membranes were stained with 0.1X CellMask™ Deep Red Plasma Membrane Stain (Invitrogen, Cat. No. C10046) at 37 °C and 150 rpm for 10 minutes. Between binding, fixation and membrane staining each three washes with DPBS were performed. The pelleted cells were resuspended in 10 µL DPBS and applied onto microscope slides (VWR, Cat. No. 651-1551).

For CbpF/CEACAM1 binding analysis to HEK293 wt cells (ATCC), 3 × 10^5^ cells per well were cultured in DMEM/F12 medium (PAN-Biotech, Cat. No. 4251184101199) supplemented with 10% (v/v) fetal bovine serum in glass bottom µ-Slide 8 Well plates (Ibidi, Cat. No. 5588389) for 2 days. Fixation was carried out with 200 µL 4% (v/v) paraformaldehyde at room temperature for 20 minutes. For binding, cells were incubated with either 1 µM AF 488 NHS-Ester conjugated CbpF nanodiscs or AF 488 conjugated nanodiscs composed of DMPC and the MSP1D1 scaffold protein without CbpF at 37 °C for 2 hours. Cell membranes were stained with 0.1X CellMask™ Deep Red Plasma Membrane Stain at 37 °C for 10 minutes. Between binding, fixation and membrane staining, three washes with DPBS were performed each. For imaging, cells were overlaid with DPBS.

Confocal images of the processed cells were acquired with an LSM710 equipped with spectral detector and PMTs (Carl Zeiss Microscopy). The LSM710 was controlled by the Zen Black software (Carl Zeiss Microscopy). For determination of protein localization, multi-color confocal imaging was performed in sequential mode with the following fluorophore-specific excitation (Ex.) and emission filter (EmF.) settings: Alexa 488 (Ex.: 488 nm; EmF.: 491–561 nm), Alexa 647 (Ex.: 633 nm; EmF.: 638–752 nm). A PL APO DIC M27 100 x/1.46 NA oil objective was used to acquire 5×5 tile stacks of seven 512×512 pixel images at a distance of 1.5 μm using a line average of 1.

Image analysis was performed using Fiji/ImageJ (Schindelin et al., 2012). Collected tiles were stitched using the Grid/collection stitching tool (Preibisch et al., 2009). The stitched images were then analyzed for integrated fluorescence of AF 488-CbpF nanodiscs using a Fiji macro provided by Dr. Tolga Soykan (Cellular imaging facility, Leibniz FMP). Fluorescence readings were then averaged and normalized for the fraction of the micrograph area covered by cells and for differences in total fluorescence intensity, providing a readout independent of laser settings and cell density.

### Surface plasmon resonance of CbpF and CEACAM1

Binding kinetics of CbpF and CEACAM1 were determined using a Biacore™ T200 surface plasmon resonance system (Cytiva). A His capture kit (Cytiva) was used to immobilize an anti-Histag antibody on a Series S Sensor Chip CM5 according to the manufacturer’s instructions. For all measurements, 25 mM MOPS-NaOH, 150 mM NaCl, 1 % P8POE(n-Octylpolyoxyethylene), pH 7.0 was used as a running buffer at a temperature of 25 °C. CEACAM1-ECD (residues 35-428) in fusion with a C-terminal hexahistidine tag (Acro Biosystems, Cat. No. CE1-H5220) was captured on the chip with a concentration of 0.5 mg/mL, a contact time of 45 sec and a flow rate of 5 µL/min, resulting in a final responsive unit (RU) value of around 35 RU. 120 µL CbpF in running buffer was injected in different concentrations ranging from 11.5 to 1480 nM, and the response difference was recorded. Analyses were performed at 25 °C with a flow rate of 30 μL/min with 240 sec for association and 600 sec for dissociation. After each run, surfaces were regenerated with 30 µL of 10 mM Glycine-HCL pH 1.5 at a flow rate of 30 μL/min for 1 min. The Biacore T200 evaluation software V.3.2.1 was used to calculate association and dissociation rate constants using a 1:1 Langmuir fitting model or a heterogeneous ligand model.

### AlphaFold2 prediction of CbpF and Cbpf/CEACAM1

A local installation of AlphaFold2 multimer (Bryant et al., 2022) was used to predict the structures of the CbpF homotrimer, which was then used as initial model in the cryo-EM map, and CbpF/CEACAM1. The sequence data base contained entries until Dec 12th, 2022. For CbpF, the full-length protein without the N-terminal signal was predicted. For CEACAM1 (Uniprot ID P13688), the full extracellular domain (residues 35-428) was predicted. Both hetero-tetrameric (3 CbpF + 1 CEACAM1) and hetero-hexameric complexes (3 CbpF + 3 CEACAM1) were predicted, and the highest-ranking models were taken for further analysis and representation.

### Complex formation of CbpF and CEACAM1

50 µL of 4.6 µM nanodisc-embedded CbpF (trimer) was incubated with 14 µM CEACAM1 ECD (Acro Biosystems, Cat no. CE1-H5220) for 60 min at 23 °C. 50 µL of the preparation were then loaded on a Superdex 200 increase 3.2-300 column equilibrated in SEC buffer using an ÄKTA pure system. Fractions of 75 µL were collected and analyzed via SDS-PAGE. The fraction with the highest total protein concentration as identified by SDS-PAGE was then used for cryo-EM.

To test CEACAM1 binding of CbpF Δ142-149, the mutant in buffer A was incubated with CEACAM1 in a 1:1 (monomer-to-monomer) molar ratio, incubated for 60 min at 23 °C, and analyzed via SEC as above. CbpF wt in buffer A with CEACAM1 served as control.

### Cryo-EM sample preparation and data acquisition of CbpF and CbpF/CEACAM1

Nanodisc-reconstituted CbpF (0.6 mg/mL total protein concentration in SEC buffer) was vitrified on freshly glow discharged Quantifoil 1.2/1.3 Cu 300 mesh grids using a Vitrobot Mark IV set to a blot force of 0, blotting time of 3.0 s, 100% humidity, and temperature of 8 ºC. The CbpF/CEACAM1 complex (0.3 mg/mL total protein concentration in SEC buffer) was vitrified under the same conditions.

Micrographs for CbpF were acquired using a FEI Titan Krios G3i microscope (TFS) operated at 300 kV equipped with a Bioquantum K3 direct electron detector and energy filter (Gatan) running in CDS super resolution counting mode at a slit width of 20 eV and at a nominal magnification of 105,000, giving a calibrated pixel size of 0.415 Å/px on the specimen level. For CbpF/CEACAM-1, an analogous setup without super resolution mode enabled was applied, giving a calibrated pixel size of 0.83 Å/px on the specimen level. EPU 2.12 (TFS) was utilized for automated data acquisition with aberration-free imaging (AFIS) enabled. For CbpF, movies were recorded for 1.2 s accumulating a total electron dose of 79.9 e^−^/Å^2^ fractionated into 30 frames. Nominal defocus values were between -1.4 and -2.6 µm. For CbpF/CEACAM1, movies were recorded for 3.2 s accumulating a total electron dose of 52.8 e^−^/Å^2^ fractionated into 60 frames, with nominal defocus values of -1.2 to -2.4 µm.

### Data processing of CbpF and CbpF/hCEACAM1-ECD complex

Cryo-EM data analysis was carried out in cryoSPARC (Punjani et al., 2017) as outlined in SI Fig. 2 (CbpF) and SI Fig. 8 (complex). For CbpF, a total of 3352 movies were acquired and then aligned using Patch Motion Correction. The CTF was then estimated using Patch CTF. Poor micrographs were manually excluded from further processing, resulting in a total of 2,930 micrographs for analysis. Particle picking was performed using the Blob Picker, resulting in 974,539 particles that were sorted using 2D classification, resulting in 260 templates. Template picking resulted in 1,387,406 particles that were extracted with a box size of 384 pixels binned to 128 pixels. Two rounds of 2D classification yielded 224,374 particles, which were re-extracted with a box size of 384 pixels without binning. After an additional round of 2D classification, 200,809 particles were used for heterogeneous refinement with enforced C3 symmetry and a refinement box size of 192 voxels. A total of 105,879 particles were assigned to the optimal initial volume, which were then used for non-uniform refinement with enforced C3 symmetry, resulting in a global resolution of 3.86 Å. Local CTF refinement and subsequent non-uniform refinement further improved the global resolution to 3.77 Å. Reference-based motion correction with subsequent local refinement produced a similarly refined map, which was then subjected to heterogeneous refinement. The best class was used for a final non-uniform refinement, resulting in a global resolution of 3.76 Å.

A total of 6,402 movies were collected for CbpF/CEACAM1 and pre-processed analogously to the CbpF dataset, resulting in a total of 6,334 micrographs for analysis. Particle picking was conducted through an iterative process of blob picking, 2D classification, and template picking. The initial blob picking of a randomly selected subset of 800 micrographs yielded 265,310 particles, which were extracted with a 384-pixel box size binned to 128 pixels. Three rounds of 2D classification yielded 72 templates, which were subsequently employed for template picking with a particle diameter of 150 Å and a minimum separation distance of 0.3 particle diameters. A total of 1,766,853 particles were extracted with a 384 pixels box size, binned to 128 pixels, and subjected to two rounds of 2D classification. The remaining 308,537 particles were re-extracted at a box size of 384 pixels without binning and employed for heterogenous refinement of a previous ab-initio reconstruction. This employed a refinement box size of 192 voxels and enforced C3 point group symmetry. A total of 211,927 particles were assigned to the optimal initial volume, which was subsequently subjected to non-uniform refinement with enforced C3 point group symmetry, resulting in a global resolution of 2.66 Å according to the gold-standard Fourier shell correlation (FSC) criterion. To enhance the quality of the resulting map, reference-based motion correction was performed, followed by another round of non-uniform refinement with enforced C3 point group symmetry, yielding a global resolution of 2.67 Å. While nominally slightly inferior in resolution to the pre-reference-based motion correction non-uniform refinement, this map was considered superior due to higher levels of visual detail.

We the aimed to increase resolution of individual CEACAM1 domains close to the periphery of the box analogous as described previously (Roderer et al., 2019). For this, initially symmetry expansion was applied, resulting in 635,781 C1-symmetric particles. These were subsequently utilized for targeted 3D classification using two distinct classes, with one bound CEACAM1 chain masked. The superior class was selected for local refinement of one CEACAM1 chain, resulting in a global resolution of 2.77 Å. Subsequently, the center of reconstruction was shifted to the most CbpF-proximal CEACAM1 domain (IgV like) using Volume Alignment Tools. The resulting 330,202 particles were re-extracted with a box size of 320 pixels without binning and reconstructed without refinement using the “Reconstruct Only” function. Subsequently, a final non-uniform refinement was conducted, with the resolved CbpF parts and one CEACAM1 chain masked, yielding a global resolution of 3.04 Å.

### Atomic modeling

Initially, the CbpF homotrimer model predicted by AlphaFold2 was rigid-body fitted into the CbpF density map that has been sharpened with DeepEMhancer (Sanchez-Garcia et al., 2021) using UCSF ChimeraX (Pettersen et al., 2021). CbpF residues Ala25 - Gly274 (3 - 252 of the mature protein) that comprise the N-terminal β roll (residues 25 - 207) were within the experimental cryo-EM density, whereas the C-terminal part of the ECD and the transmembrane region were not resolved. The latter parts were removed and the model was then manually adjusted in Coot (WinCoot version 0.9.8.7) (Emsley et al., 2010), followed by real-space refinement in PHENIX (Afonine et al., 2018).

For the CbpF/CEACAM1 complex, the CbpF model from above and three AlphaFold2 predicted models of the extracellular part of CEACAM1 obtained from UniProt (accession number P13688) were fitted in the density map of the complex using UCSF ChimeraX. For the latter, the IgC2-like domains B and A2 were not resolved and removed, whereas the N-terminal IgV-like domain (residues 35-143) and the IgC2-like A1 domain (residues 144-234) were fitted individually and re-connected in Coot. The models were then manually adjusted in Coot, where glycans (N-acetylglucosamines) were attached to non-modeled densities of CEACAM1, and the models were refined using real-space refinement in PHENIX. Modeling of CEACAM1 was carried out in the density subtracted and locally refined map (SI Fig. 7I-N) and the resulting model was then placed in the full map (SI Fig. 7G-L).

Statistics of the cryo-EM data and the refined atomic models are summarized in SI Table 1.

## Data availability

The models and cryo-EM maps of CbpF, CbpF/CEACAM1 complex, and CEACAM1 in symmetry-expanded and shifted complex have been deposited in the PDB and EMDB under accession numbers 9GH4 and 51346, 9GH5 and 51347, and 9GH6 and 51348, respectively.

## Acknowledgements

We acknowledge access to electron microscopic equipment at the Core Facility for cryo-Electron Microscopy (CFcryoEM) of the Charité - Universitätsmedizin Berlin supported by DFG (INST 335/588-1 FUGG) for cryo-EM data collection, and we thank Dr. Thiemo Sprink for recording the data. We gratefully acknowledge access to and training for confocal microscopes provided by the Cellular Imaging Facility at the Leibniz FMP (headed by Dr. Martin Lehmann), and the help of Dr. Tolga Soykan in quantitative evaluation of the cellular images. We acknowledge mass spectrometric analysis carried out by Heike Stephanowitz and Max Ruwolt (Mass Spectrometry Core facility, Leibniz FMP).

