## Supplementary information for "Structural basis for immune cell binding of *Fusobacterium nucleatum* via the trimeric autotransporter adhesin CbpF"

Marongiu G.L. et al.

Supplementary Figures

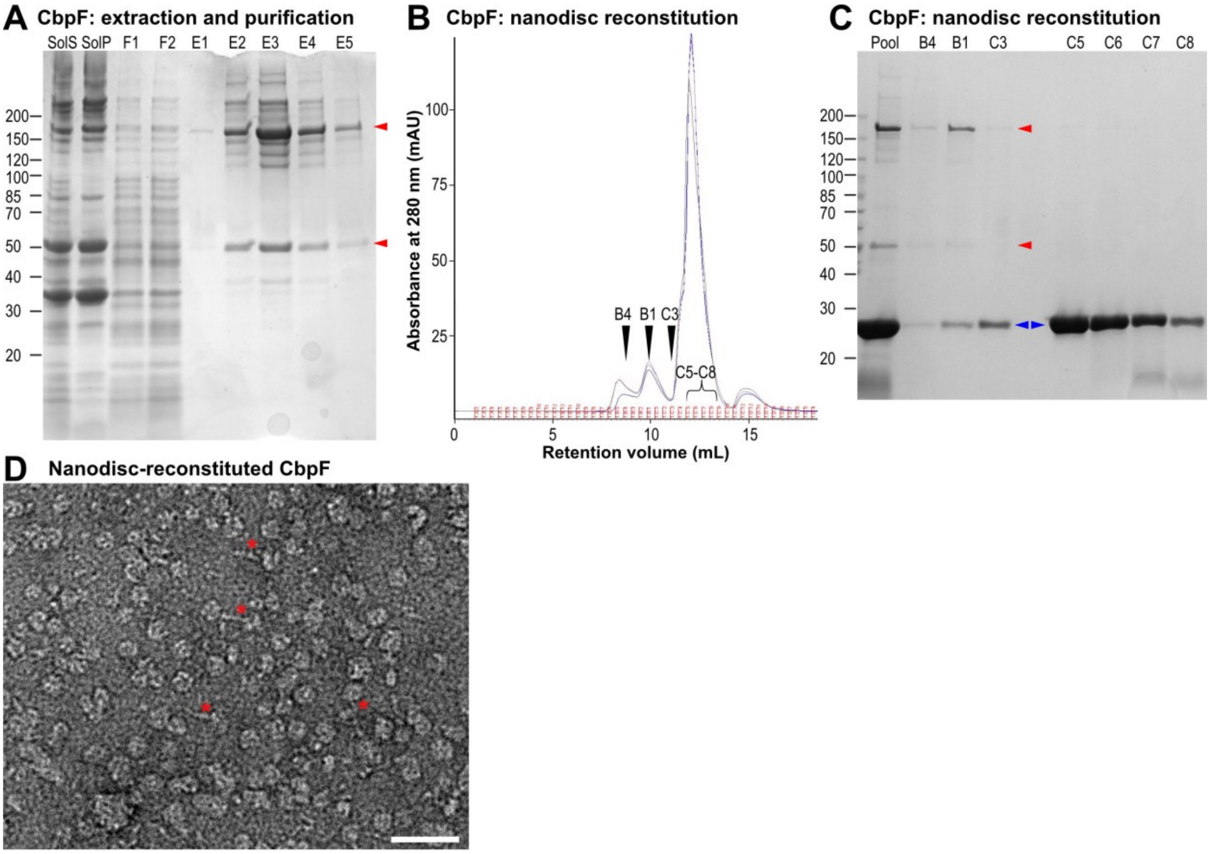


**Supplementary Figure 1: Purification and nanodisc reconstitution of CbpF from *F. nucleatum* ATCC25586.** A: SDS-PAGE showing the purification of memberane-extracted CbpF using StrepTactin resin. SolS/SolP: supernatant and pellet after solubilization, F1/F2: flowthrough fractions 1/2, E1-E5: elution fractions. B: Chromatogram of CbpF after nanodisc reconstitution (CbpF-to-MSP1D1-to-lipid ratio of 1:5:80) on a Superdex 200 increase 10/300 column. The three chromatograms show three consecutive runs of a reconstituted sample, where 500 µL pool were loaded each. The fractions shown in C are indicated. C: SDS-PAGE showing nanodisc reconstitution of CbpF and purification via size exclusion chromatography. The fractions on the gel are indicated in B. D: Section of a negative stain EM micrograph of nanodisc-reconstituted CbpF (fraction B1 of B,C), recorded at 120 kV. The red asterisks indicate rod-shaped CbpF that protrudes from nanodiscs. Scale bar: 50 nm.

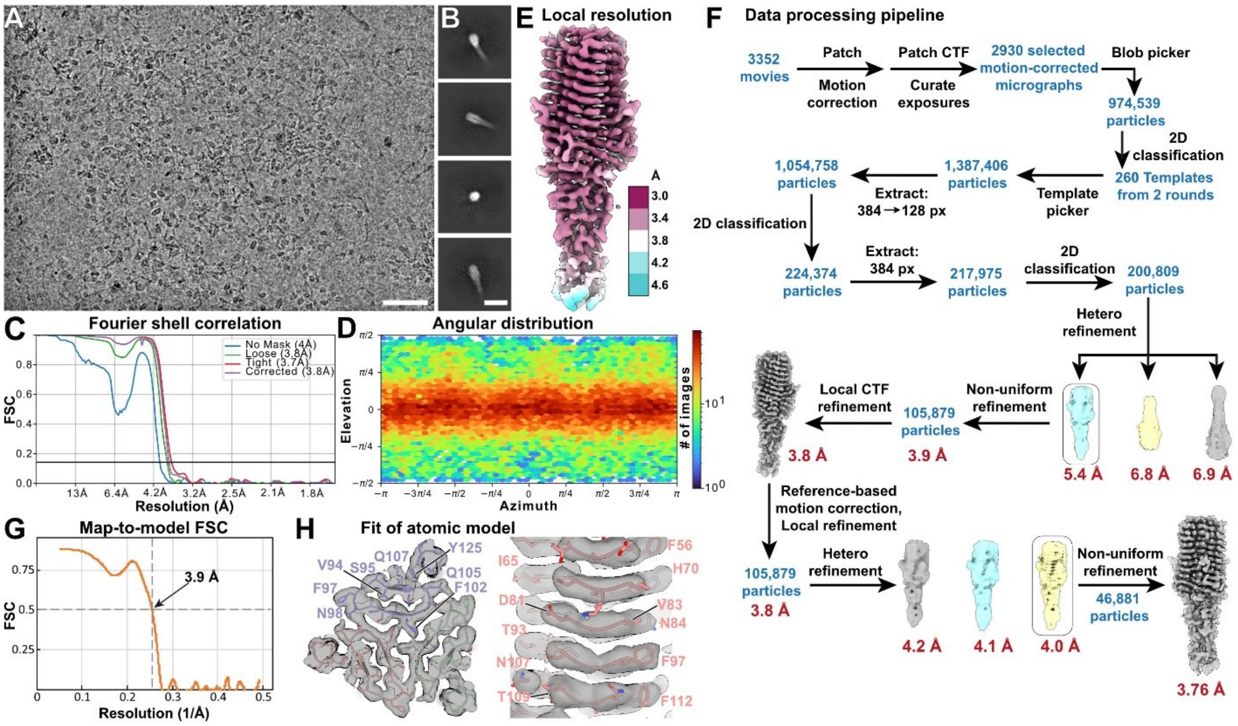


**Supplementary Figure 2: Cryo-EM and SPA of CbpF.** A: Representative cryo-EM micrograph recorded at 300 kV and -1.8 µm defocus. Scale bar: 50 nm. B: Representative 2D class averages, showing the membrane-distal domain of CbpF in side views and top view. Scale bar: 10 nm. C,D,E: Fourier shell correlation (C), angular distribution (D), and density map colored by local resolution (E) of the final cryo-EM density map of CbpF from 46,881 particles with C3 symmetry applied. F: SPA data processing scheme applied for CbpF. All steps were carried out in cryoSPARC, and all refinements after the first Hetero refinement were carried out with C3 symmetry. G: Map-to-model correlation of the CbpF model (residues 3-252) against the final non-sharpened density map. H: Fit of atomic model in selected parts of the density map. Residues for one of the three protomers are indicated. The density maps in E,F and H have been sharpened with DeepEMhancer.

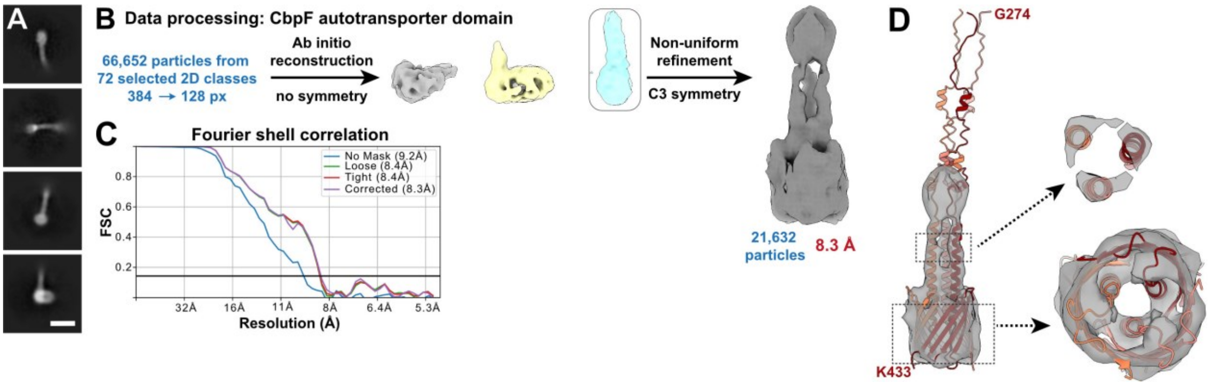


**Supplementary Figure 3: The autotransporter domain of CbpF.** A: Representative 2D class averages that show nanodisc density with protruding CbpF. The class averages originate from the cryo-EM data as in SI Figure 2. Scale bar: 10 nm. B: Data processing of particles underlying the 2D classes that show nanodisc-embedded CbpF autotransporter domains. C: Fourier shell correlation of the density map of the CbpF autotransporter domain, originating from 21,632 particles and C3 symmetry. D: Fit of the AlphaFold2 prediction of CbpF (residues 274 - 433) to the autotransporter density map. The three protomers were fitted independently.

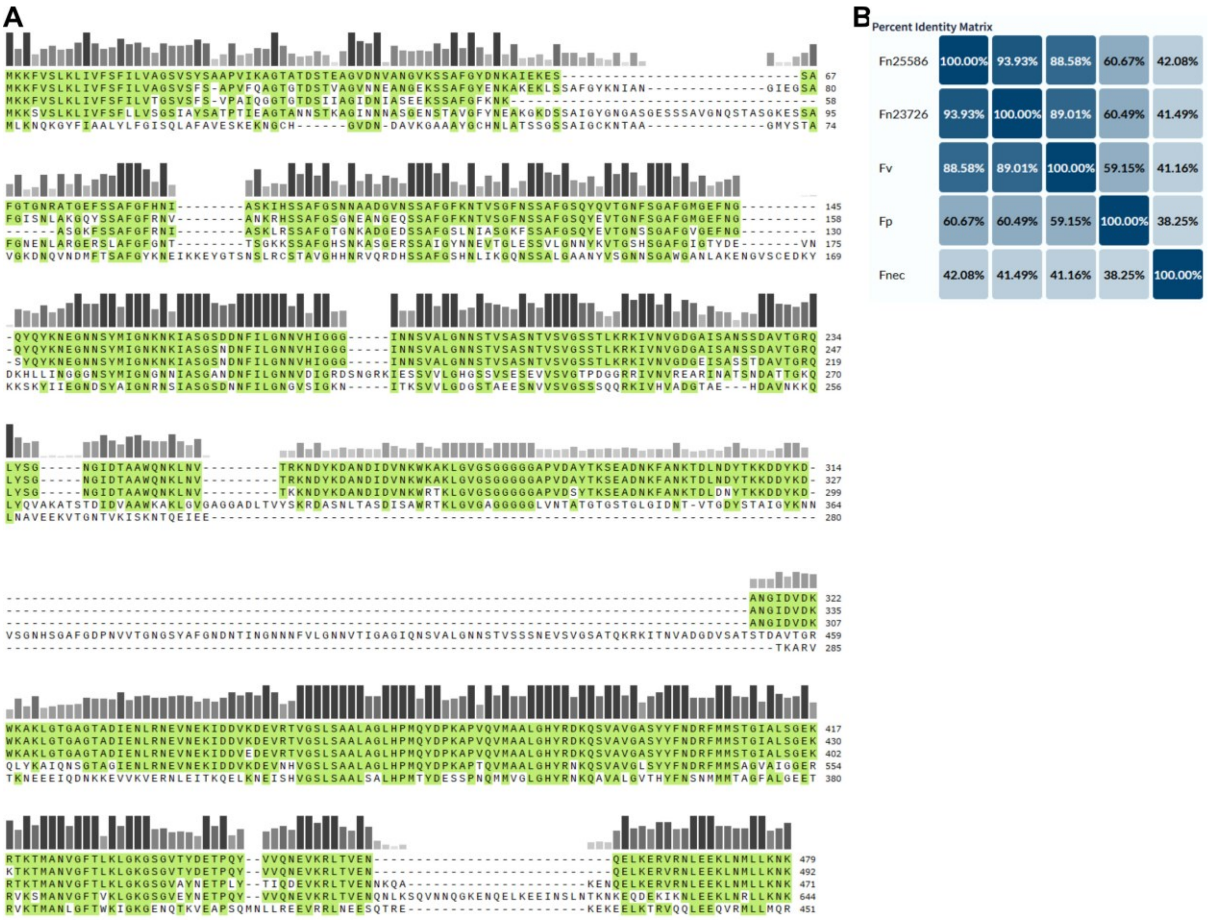


**Supplementary Figure 4: Comparison of CbpF in *Fusobacterium nucleatum* strains ATCC25586, ATCC23726, *F. vincentii*, *F. polymorphum*, and *F. necrophorum*.** A: Full sequence alignment from which the excerpt in Fig. 2A is derived. The order of sequences from top to bottom is as indicated above. B: Identity matrix of the five Fn sequences.

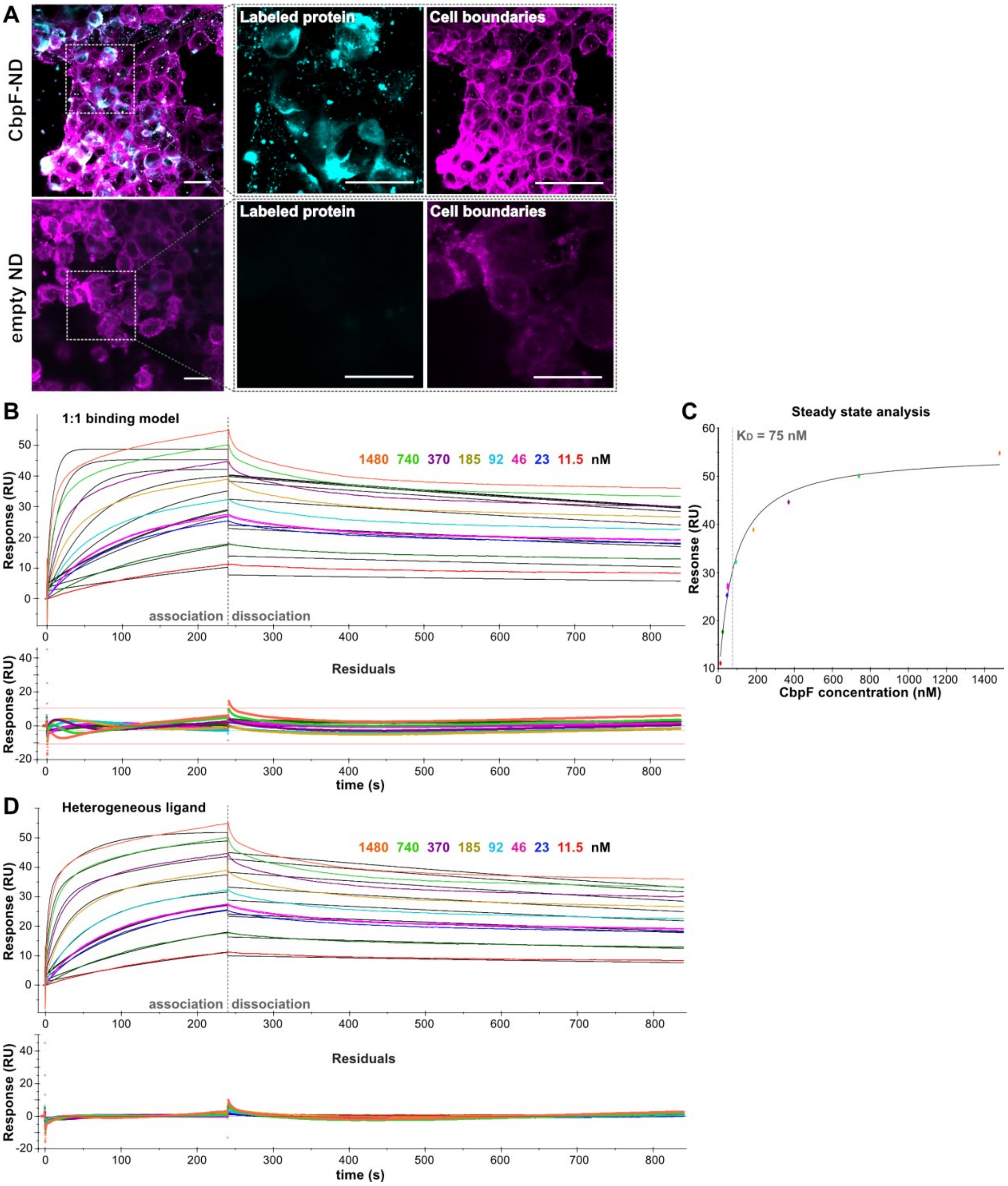


**Supplementary Figure 5: Cell binding of CbpF and SPR of CbpF with CEACAM1.** A: Representative micrographs that illustrate binding of CbpF to human embryonic kidney (HEK) cells, whereas empty nanodiscs do not bind. B: SPR sensorgram of CbpF and immobilized CEACAM1-ECD, as in Fig. 3C, with the concentrations of CbpF indicated. The black solid lines show a global fit of the binding curves according to a 1:1 binding model. The lower panel shows the residuals of the fit. C: Steady-state affinity of the SPR data as shown in Fig. 3C. The fit revealed a $K_D$ of 75 ± 14 nM. D: SPR sensorgram as in C with the data fitted according to a heterogeneous ligand model. $K_D$ values of 2.5 and 47 nM were obtained.

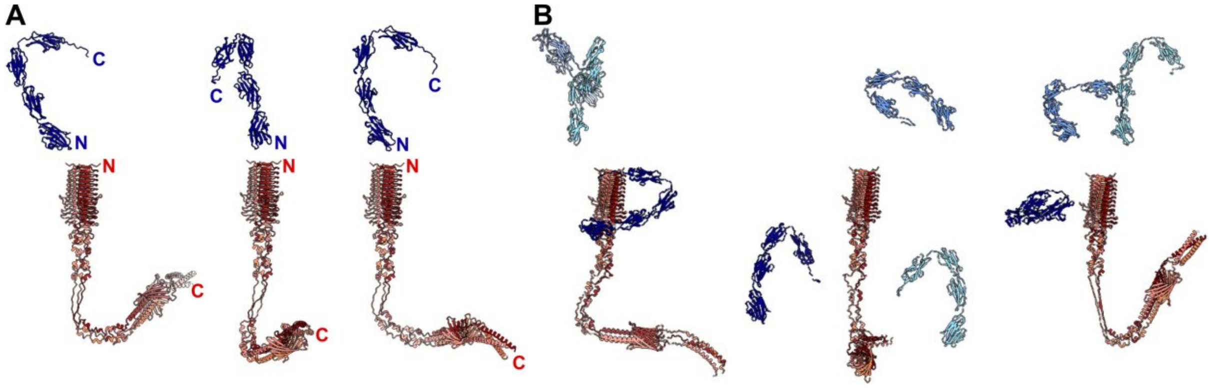


**Supplementary Figure 6: Representative AlphaFold2 complex predictions of CbpF (shades of red) and CEACAM1 (shades of blue).** A: Three highest-rated predictions of 3:1 complex. N- and C-termini of CEACAM1 and one CbpF within the trimer are indicated. B: Three highest-rated predictions of 3:3 complex. No meaningful and reproducible CbpF/CEACAM1 complexes could be predicted in both cases.

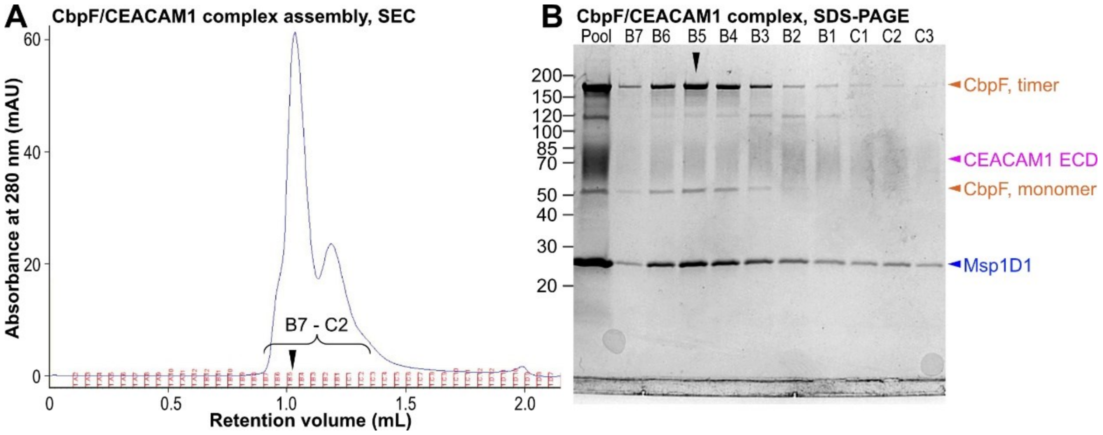


**Supplementary Figure 7: Assembly of nanodisc-reconstituted CbpF with CEACAM1-ECD.** A: Chromatogram of CbpF/CEACAM1 (1:1 molar ratio) on a Superdex 200 increase 3.2/300 column. The fractions shown in B are indicated. B: SDS-PAGE showing fractions of SEC as in A. Bands corresponding to CbpF, CEACAM1 and the nanodisc scaffold protein MSP1D1 are indicated by orange, magenta, and blue arrowheads. The fraction B5 used for cryo-EM is indicated by a black arrowhead in A and B.

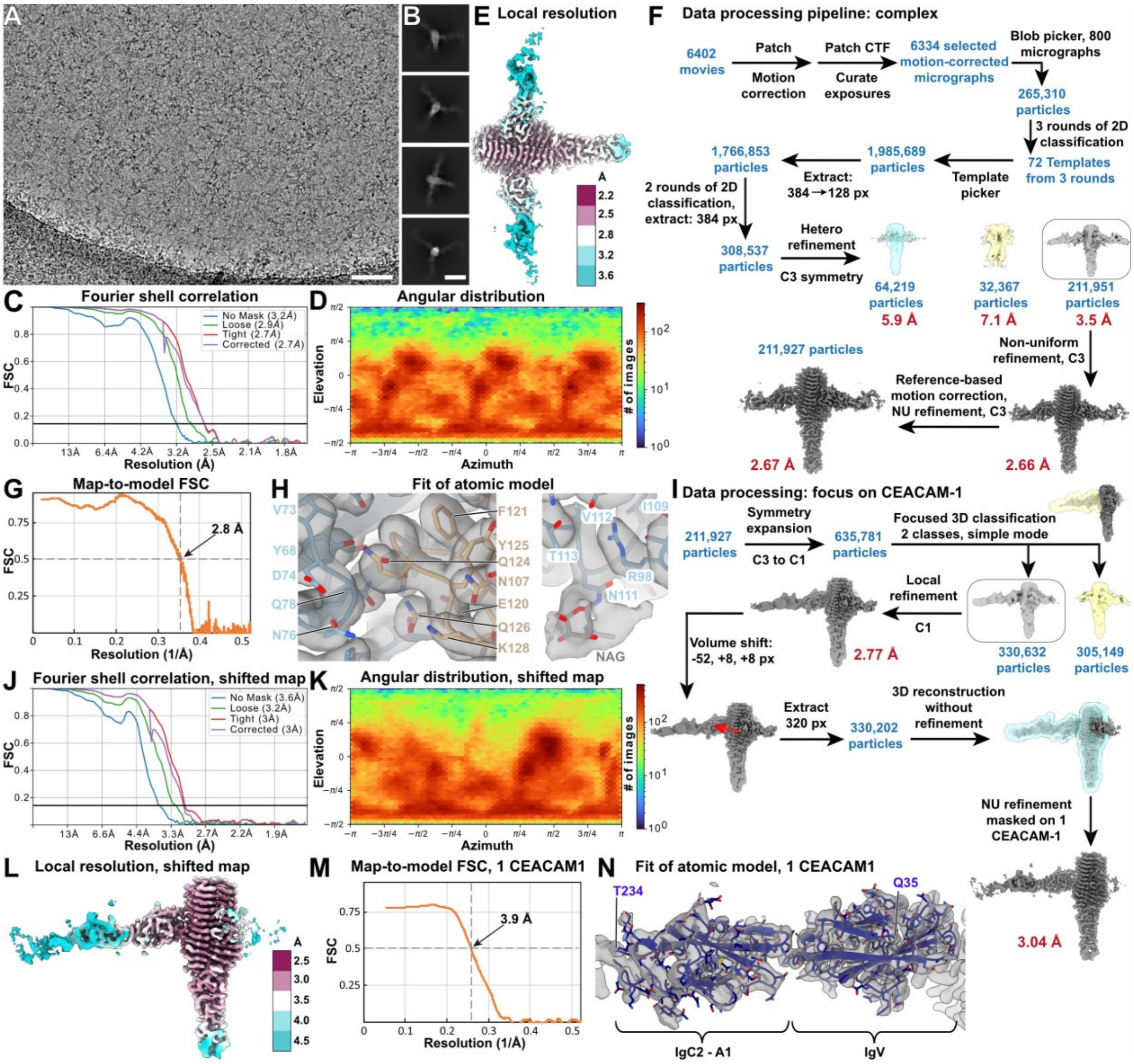


**Supplementary Figure 8: Cryo-EM and SPA of CbpF/CEACAM1 complex.** A: Representative cryo-EM micrograph recorded at 300 kV and -1.5 µm defocus. Scale bar: 50 nm. B: Representative 2D class averages, showing different views of the complex. Scale bar: 10 nm. C,D,E: Fourier shell correlation (C), angular distribution (D), and density map colored by local resolution (E) of the final cryo-EM density map of the complex from 211,927 particles with C3 symmetry applied. F: SPA data processing scheme applied for the complex. All steps were carried out in cryoSPARC, and all refinements were carried out with C3 symmetry. G: Map-to-model correlation of the CbpF/CEACAM1 model (heterotetramer; CbpF residues 4 - 252, CEACAM1 residues 35 - 234) against the final non-sharpened density map. H: Fit of atomic model in selected parts of the density map. Left: CbpF/CEACAM1 interface, right: selected glycosylation of CEACAM1. I: Data processing scheme with the center of reconstruction shifted towards one CEACAM1. The red arrow indicates the shifting direction and distance. J,K,L: Fourier shell correlation (J), angular distribution (K), and density map colored by local resolution (L) of the cryo-EM density map of the complex with center of reconstruction shifted to one CEACAM1, originating from 330,202 particles without symmetry. M: Map-to-model correlation of one CEACAM1 fitted (residues 35 - 234) into the shifted density map. N: Fit of atomic model as of M. The density maps in E,F,H, I, L and N have been sharpened with DeepEMhancer.

### Supplementary Table

Supplementary Table 1: Cryo-EM data collection, refinement and validation statistics

| | CbpF<br>PDB 9GH4 | CbpF/CEACAM1<br>PDB 9GH5 | 1 CEACAM1 in complex<br>PDB 9GH6 |
|---|---|---|---|
| **Data collection and processing** | | | |
| Magnification | | 105,000 | |
| Voltage (kV) | | 300 | |
| Camera | | Gatan K3 with energy filter | |
| Electron exposure (e–/Å$^2$) | 79.9 | 52.8 | |
| Defocus range (μm) | -1.4 – 2.6 | -1.2 – 2.4 | |
| Pixel size (Å) | 0.83 (0.415 super res) | 0.83 | |
| Micrographs used | 2930 | 6,334 | |
| Total extracted particle images | 1,054,758 | 1,766,853 | |
| Refined particle images | 200,809 | 308,537 | 635,781 |
| Final particle images | 46,881 | 211,927 | 330,202 |
| Map resolution (Å) | 3.76 | 2.67 | 3.04 |
| at FSC threshold | 0.143 | 0.143 | 0.143 |
| Map resolution range (Å) | 3.0 – 4.6 | 2.2 – 3.6 | 2.5 – 4.5 |
| **Refinement** | | | |
| Refinement package | | Phenix 1.21-5207 | |
| Model resolution (Å) | 3.9 | 2.8 | 3.9 |
| at FSC threshold | 0.5 | 0.5 | 0.5 |
| Map sharpening *B* factor (Å$^2$) | -132.6 | -115.8 | -132.8 |
| Model composition | | | |
| Non-hydrogen atoms | 5454 | 10,383 | 1661 |
| Protein residues | 750 | 1347 | 200 |
| Ligand | 0 | NAG: 18 | NAG: 7 |
| Water | 0 | 3 | 0 |
| *B* factors (min/max/mean, Å$^2$) | | | |
| Protein | 75.16/167.36/105.25 | 65.23/370.39/147.09 | 125.60/284.48/191.15 |
| Ligand | | 111.51/359.89/248.44 | 151.79/278.25/235.50 |
| Water | | 77.35/82.83/79.88 | |
| R.m.s. deviations | | | |
| Bond lengths (Å) | 0.002 | 0.003 | 0.003 |
| Bond angles (°) | 0.599 | 0.715 | 0.607 |
| Validation | | | |
| MolProbity score | 1.53 | 2.08 | 1.77 |
| Clashscore | 7.84 | 9.98 | 8.31 |
| Poor rotamers (%) | 1.06 | 2.54 | 1.12 |
| Ramachandran plot | | | |
| Favored (%) | 97.58 | 96.25 | 95.96 |
| Allowed (%) | 2.42 | 3.75 | 4.04 |
| Disallowed (%) | 0.00 | 0.00 | 0.00 |

**Supplementary movie legend**

Supplementary Movie 1: Illustration of the structure of the CbpF/CEACAM1 complex, highlighting one binding interface. The CbpF homotrimer is shown in red tones, and three bound CEACAM1 molecules are shown in blue tones.
